# Evolution of selenophosphate synthetases: emergence and relocation of function through independent duplications and recurrent subfunctionalization

**DOI:** 10.1101/014928

**Authors:** Marco Mariotti, Didac Santesmasses, Salvador Capella-Gutierrez, Andrea Mateo, Carme Arnan, Rory Johnson, Salvatore D’Aniello, Sun Hee Yim, Vadim N Gladyshev, Florenci Serras, Montserrat Corominas, Toni Gabaldón, Roderic Guigé

## Abstract

SPS catalyzes the synthesis of selenophosphate, the selenium donor for the synthesis of the amino acid selenocysteine (Sec), incorporated in selenoproteins in response to the UGA codon. SPS is unique among proteins of the selenoprotein biosynthesis machinery in that it is, in many species, a selenoprotein itself, although, as in all selenoproteins, Sec is often replaced by cysteine (Cys). In metazoan genomes we found, however, SPS genes with lineage specific substitutions other than Sec or Cys. Our results show that these non-Sec, non-Cys SPS genes originated through a number of independent gene duplications of diverse molecular origin from an ancestral selenoprotein SPS gene. Although of independent origin, complementation assays in fly mutants show that these genes share a common function, which most likely emerged in the ancestral metazoan gene. This function appears to be unrelated to selenophosphate synthesis, since all genomes encoding selenoproteins contain Sec or Cys SPS genes (SPS2), but those containing only non-Sec, non-Cys SPS genes (SPS1) do not encode selenoproteins. Thus, in SPS genes, through parallel duplications and subsequent convergent subfunctionalization, two functions initially carried by a single gene are recurrently segregated at two different loci. RNA structures enhancing the readthrough of the Sec-UGA codon in SPS genes, which may be traced back to prokaryotes, played a key role in this process. The SPS evolutionary history in metazoans constitute a remarkable example of the emergence and evolution of gene function. We have been able to trace this history with unusual detail thanks to the singular feature of SPS genes, wherein the amino acid at a single site determines protein function, and, ultimately, the evolutionary fate of an entire class of genes.

## Introduction

Selenoproteins are proteins that incorporate the non standard amino acid selenocysteine (Sec or U) in response to the UGA codon. The recoding of UGA, normally a stop codon, to code for Sec is arguably the most outstanding programmed exception to the genetic code. Selenoproteins are found, albeit in small numbers, in organisms across the entire tree of life. Recoding of UGA is mediated by RNA structures within selenoprotein transcripts, the SECIS (SElenoCysteine Insertion Sequence) elements. Sec biosynthesis and insertion also require a dedicated system of trans-acting factors, called the Sec machinery, including elements that are common and others that are specific to the three domains of life: bacteria (Kryukov and Gladyshev 2004)(Yoshizawa and Bock 2009), archaea (Rother et al. 2001), and eukaryotes (Squires and Berry 2008)(Allmang et al. 2009).

The very existence of selenoproteins is puzzling. Sec can apparently be substituted by cysteine (Cys)—as often happens during evolution— with seemingly a small or null impact on protein function. In fact, selenoproteins may be absent in an entire taxonomic group, but present in sister lineages. This can be seen most dramatically within fruit flies: while *Drosophila melanogaster* and most other flies possess three selenoprotein genes, their relative *Drosophila willistoni* has completely replaced Sec with Cys in proteins and lost the capacity to synthesize selenoproteins (Chapple and Guigó, 2008)(Lobanov et al. 2008). Fungi and plants also lost this capacity (Lobanov et al. 2009). In other cases, however, such as in *Caenorhabditis elegans*, the entire pathway is maintained only to synthesize a single selenoprotein (Taskov et al. 2005). It appears that selective advantage exists to maintain Sec, at least in vertebrates, since strong purifying selection across Sec sites against mutations to Cys has been reported in this lineage (Castellano et al. 2009). Sec encoding has been hypothesized to be an ancestral trait, already present in the early genetic code, since a number of selenoprotein families are shared between prokaryotes and eukaryotes. However, the evolutionary continuity of the Sec recoding systems across domains of life is not certain, since it would require the translocation of the SECIS element (within the coding region in bacteria but within the 3’ UTR in eukaryotes) and also a radical alteration of its structure, for both of which it is difficult to postulate plausible evolutionary scenarios.

Selenophosphate synthetase (SPS, also called SelD or selenide water dikinase) is unique among the components of the Sec machinery in that it is often a selenoprotein itself. SPS catalyzes the synthesis of selenophosphate (SeP) from selenide, ATP and water, producing AMP and inorganic phosphate as products. SeP is the selenium (Se) donor for the synthesis of Sec, which, in contrast to other amino acids, takes place on its own tRNA (Xu et al. 2007a)(Palioura et al. 2009).

SPS proteins are conserved from bacteria to human with about 30% identity and are found in all species known to encode selenoproteins. In prokaryotes, SPS (SelD) is found also in species where SeP is used to produce selenouridine (SeU) in tRNAs, where it acts as the Se donor to protein ybbB (selenouridine synthase). The presence of the two traits (SeU and Sec) overlaps, but not completely (Romero et al. 2005). In eukaryotes, SPS is generally found as a selenoprotein (SPS2), while in prokaryotes homologues with Cys aligned to the Sec position are common. Evolutionary conversion of a Sec residue to Cys is a common process and it has been reported within prokaryotes (Zhang et al. 2006), insects (Chapple and Guigó 2008), and vertebrates (Mariotti et al. 2012). Cys homologues of SPS conserve molecular function, although catalytic efficiency or substrate specificity may change. For instance, substitution of Sec to Cys decreased (but did not abolish) SPS2 activity in mouse (Kim et al. 1997).

In vertebrates and insects, two paralogous SPS genes have been reported: SPS2, which is a selenoprotein, and SPS1, which is not, and carries a threonine (Thr) in vertebrates and an arginine (Arg) in insects in place of Sec (Xu et al. 2007b). In contrast to Cys conversion, Thr or Arg conversion seems to have significantly altered the molecular function of the protein. While SPS2 has been shown to produce SeP, SPS1 is inactive in this reaction. Murine SPS1 does not generate SeP *in vitro*, and does not even consume ATP in a Se dependent way (Xu et al. 2007a). Consistently, knockout of SPS1 in mouse cell lines has been shown not to affect selenoprotein synthesis (Xu et al. 2007b). *Drosophila* SPS1 too was shown to lack the ability to catalyze selenide-dependent ATP hydrolysis or to complement SPS deficiency in *Escherichia coli* (Persson et al. 1997). In insects, SPS1 is preserved in species that lost selenoproteins (Chapple and Guigó 2008). Although human SPS1 interacts with Sec synthase (SecS) (Small-Howard et al. 2006), these findings, taken as a whole, suggest that SPS1 functions in a pathway unrelated to selenoprotein biosynthesis (Lobanov et al. 2008). What the function of SPS1 may be remains an open question. Human SPS1 has been proposed to function in Sec recycling, since a *E. coli* SelD mutant can be rescued by SPS1 but only when grown in the presence of L-selenocysteine (Tamura et al. 2004). In *Drosophila*, SPS1 has been proposed to be involved in vitamin B6 metabolism (Lee et al. 2011) and in redox homeostasis, since it protects from ROS induced damage (Morey et al. 2003).

Here we study the evolutionary history of SPS genes across the entire tree of life. We found that the presence of Sec/Cys SPS genes, together with a few other gene markers, recapitulates well the Se utilization traits (Sec and SeU) in prokaryotic genomes. Within eukaryotes, specifically within metazoans, we detected a number of SPS homologues with amino acids other than Sec or Cys at the homologous UGA position. We found that Cys- or Sec-containing SPS genes (SPS2) are found in all genomes encoding selenoproteins, while genomes that contain only SPS genes carrying amino acids other than Sec or Cys at the homologous UGA position (SPS1) do not encode selenoproteins. In SPS genes, thus, the amino acid occurring at a single position is unequivocally associated to protein function, and to the evolutionary fate of an entire class of proteins. This feature, that may be unique among all protein families, makes SPS genes singularly appropriate to investigate evolution of gene function. Indeed, thanks to this feature, we have been able to untangle the complex history of SPS genes with exceptional detail. Our analysis reveals that SPS1 genes in different metazoan lineages (including those of human and fly) originated from an ancestral Sec-carrying SPS gene through duplication events in parallel lineages of metazoans. Despite originated independently, SPS1 genes share a common function, and this likely emerged in the ancestral metazoan parental gene. This indicates selective pressure during metazoan history to segregate different functions to separate loci, and constitutes a remarkable example of recurrent escape from adaptive conflict through gene duplication and subfunctionalization (Hittinger and Carroll, 2007). Within insects, after SPS duplication, a number of taxa lost the Sec encoding SPS2 gene, and therefore the capacity to synthesize selenoproteins (Chapple and Guigo, 2008)(Lobanov et al. 2008)—becoming, together with some nematodes (Otero et al. 2014), the only known selenoproteinless metazoans. Strikingly, SPS1 in selenoproteinless hymenoptera conserved the ancestral UGA codon. Our analyses point out that UGA readthrough in hymenopterans is enhanced by overlapping RNA structures, also present in other selenoproteins, that resembles that of the bacterial SECIS, uncovering a possible evolutionary link between the prokaryotic and eukaryotic Sec encoding systems. This family of readthrough-enhancing RNA elements would have played a key role throughout all the evolution of SPS genes, and particularly in the emergence of the SPS1 function in the metazoan ancestral Sec SPS gene.

## Results

Selenoprotein genes are usually miss-annotated in prokaryotic and eukaryotic genomes, due to the recoding of UGA, normally a stop codon, to Sec. Therefore, we used Selenoprofiles (Mariotti and Guigo 2010), a dedicated selenoprotein prediction tool, to search for SelD/SPS genes in all fully-sequenced eukaryotic and prokaryotic genomes (505 and 8,263, respectively). We then utilized a combination of approaches to reconstruct their phylogenetic history. Methods and analyses are fully discussed in Supplementary Material S1-S5.

### SelD as a marker for selenium utilization in prokaryotes

Figure 1 shows the distribution of SelD (i.e. prokaryotic SPS) genes in a reference set of 223 prokaryotic genomes (Pruitt et al. 2012), along with the presence of other Se utilization gene markers. The occurrence of SelD and other Sec machinery components is in good agreement with previous findings (Zhang et al. 2006)(Zhang et al. 2008)(Zhang and Gladyshev 2008)(Zhang et al. 2010). Supplementary Material S1 contains details of the genes found in each major lineage investigated, both in the reference set, and in the extended set of 8,263 genomes.

**Figure 1:**
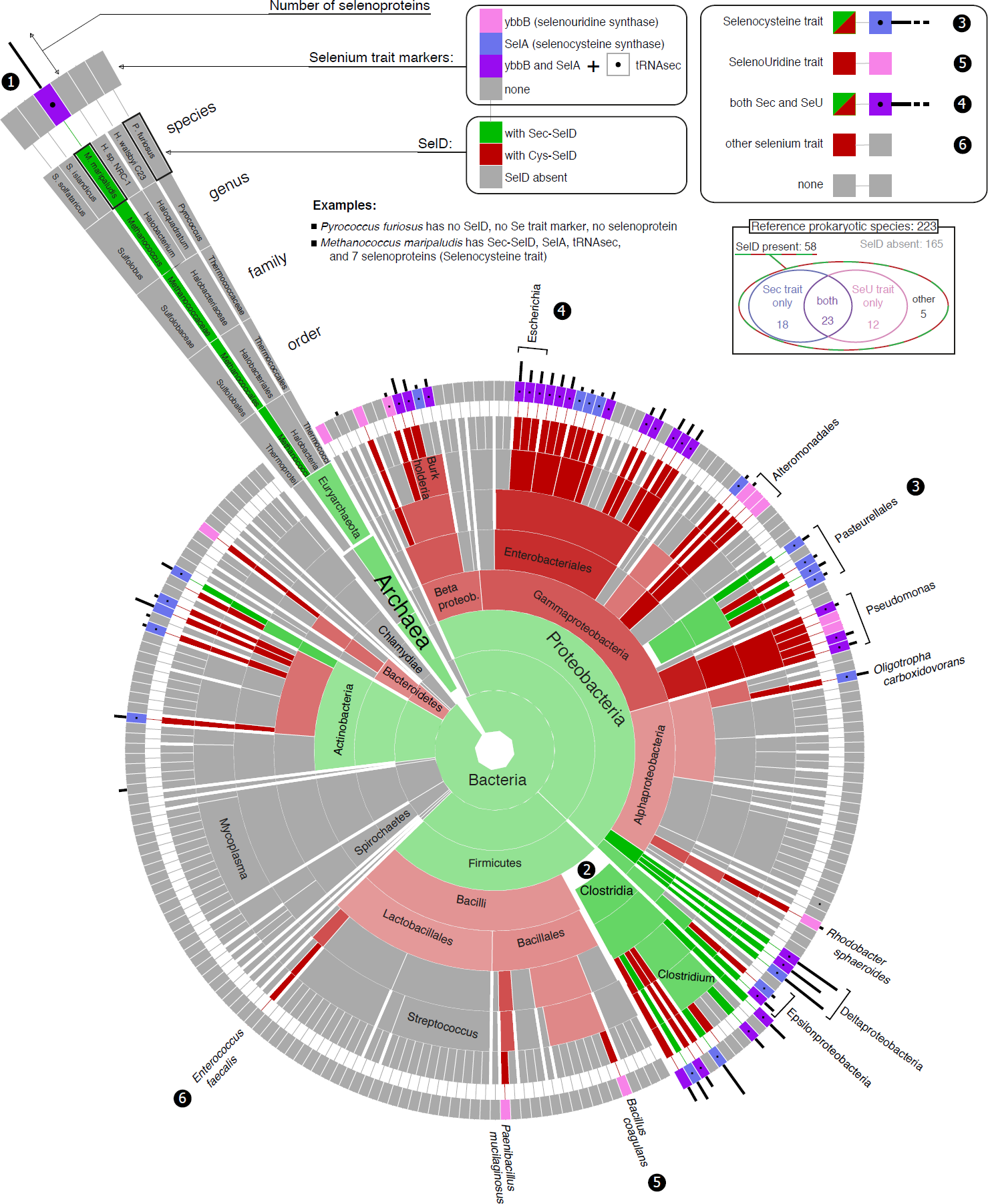
Phylogenetic profile of SPS and Se utilization traits in prokaryotes. The sunburst tree shows the phylogenetic structure of the reference set of 223 prokaryotic genomes (taken from NCBI taxonomy), and the presence of SelD genes and other markers of Se utilization. The section for archaea is zoomed in as a guide to interpret the plot (1). Every ring-shaped section represents a taxonomic rank in NCBI taxonomy (superkingdom, phylum, class, order, family, genus). The last two outermost rings display features for each of the species analyzed. The length of the black bars protruding from the outermost circle is proportional to the number of selenoproteins detected in each species. The outermost ring is color coded for the presence of ybbB and SelA in the species, with a black dot inside for denoting tRNAsec presence. The second outermost ring is labeled for SelD presence and type: “SelD-Sec”, “SelD-Cys”, “no gene found”. The color is propagated to the lower ranks by hierarchy. Transparency is used to display how many species under a lineage have the same label. Assuming no Cys to Sec conversion and no horizontal transfer, colors reflect the predicted SelD presence at ancestral nodes. This allows detecting the Sec to Cys conversions manually, for example in Clostridia (2). The hierarchical color assignment is violated only in the case of *Gammaproteobacteria*, altered to be red. In fact, although its sublineage *Pasteurellales* contains SelD-Sec (see 3), our analysis points to a Cys to Sec conversion instead, which implies that the ancestral state for Gammaproteobacteria was SelD-Cys. The plot can be used to map the Sec and selenouridine (SeU) traits (see top-right panel). For example *Escherichia coli* (4) has both traits, Pasteurellales only Sec (3), and *Bacillus coagulans* only SeU (5). *Enterococcus faecalis* has a Cys-SelD gene, but no other Se utilization marker (6) (Romero et al. 2005)(Zhang et al. 2008). Expanded versions of the plot (up to 8,263 species) are available in Supplementary Material S1. Note that gene fusions and extensions are not represented in this plot.

SelD genes were found in 26% of the reference prokaryotic genomes. A significant fraction (19%) of the detected SelD genes encoded a protein with a Sec residue (always in the same position), with all the rest containing Cys instead. The occurrence of SelD is, in general, consistent with that of the rest of the machinery for Sec (SelA, tRNAsec) and/or for SeU (ybbB), and also with selenoprotein presence (see Figure 1). The Sec trait (SelD, SelA, tRNAsec, selenoproteins) was found in a slightly larger group of organisms than the SeU trait (SelD, ybbB): 18% vs 16%, respectively. The two traits showed a highly significant overlap (p-value < 0.0001, one-tailed Fisher’s exact test): 66% of species with the SeU trait also had the Sec trait, 56% of species with Sec also had SeU, and 10% of all species had both. The Sec and SeU markers showed a scattered distribution across the prokaryotic tree, reflecting the dynamic evolution of Se utilization. The complexity of this phylogenetic pattern is even more evident when considering the extended set of prokaryotic species (see Supplementary Material S1 and Figure SM1.1).

In almost every species (93%) with SelD, SelA and/or ybbB were identified, indicating the utilization of SeP for Sec and/or SeU. A notable exception was the *Enterococcus* genus, where many species including *Enterococcus faecalis* possessed SelD, but no other markers of Se usage. This had already been reported as an indicator of a potential third Se utilization trait (Romero et al. 2005)(Zhang et al. 2008). Se is in fact used by these species as a cofactor to molybdenum hydroxylases (Haft and Self 2008)(Srivastava et al. 2011).

Among archaea, SelD was found only in *Methanococcales* and *Methanopyri* genomes, whose selenoproteins have been previously characterized (Stock and Rother 2009). The SeU trait was found only in *Methanococcales*, although with a peculiarity: ybbB is split in two adjacent genes (Su et al. 2012). In *Pasteurellales*, an Order within *Gammaproteobacteria*, we identified a bona-fide Cys to Sec conversion. Most of *Gammaproteobacteria* possess a SelD-Cys (or none), and Sec forms are found almost uniquely in *Pasteurellales*. Phylogenetic sequence signal supports codon conversion rather than horizontal transfer as the cause for SelD-Sec (Supplementary Material S1). Even though cases of conversion of Cys to Sec have been proposed (Zhang et al. 2006), this is the first clearly documented case.

Previous reports have described some SPS2 genes fused to other genes (Zhang et al. 2008)(da Silva et al. 2013). Thus, we used a computational strategy to identify gene fusions or extensions in our SelD gene dataset (see Methods and Supplementary Material S2). Fusions with a NADH dehydrogenase–like domain (Zhang et al. 2008) are by far the most common, and they are found scattered across a wide range of bacteria (Figure SM2.1). We also detected two instances of fusions with the NifS-like protein (Cys sulfinate desulfinase, proteins that deliver Se for the synthesis of SeP by SPS2, (Lacourciere et al. 2000)). In all cases, the extension/fusion is on the N-terminal side of the SPS2 genes, and these are always SelD-Cys with the single exception of NifS-SPS in *Geobacter sp. FRC-32*, which contains Sec. Since we found selenoproteins and other Sec machinery genes in all these genomes, we predict that generally these extended SPS genes have retained the original SeP biosynthetic activity.

### SPS2 as a marker for Sec utilization in eukaryotes

Figure 2 shows SPS genes and predicted selenoproteins found in a representative set of eukaryotic genomes. The presence of SPS2 genes (defined as those with Sec, or Cys) correlates perfectly with the presence of selenoproteins. Thus, SPS2 is a marker of Sec utilization in eukaryotes. Our results replicate and substantially expand to unprecedented coverage previous surveys of selenoproteins in eukaryotic genomes (e.g. (Lobanov et al. 2007)(Chapple and Guigó 2008)(Lobanov et al. 2009)(Jiang et al. 2012)).

**Figure 2:**
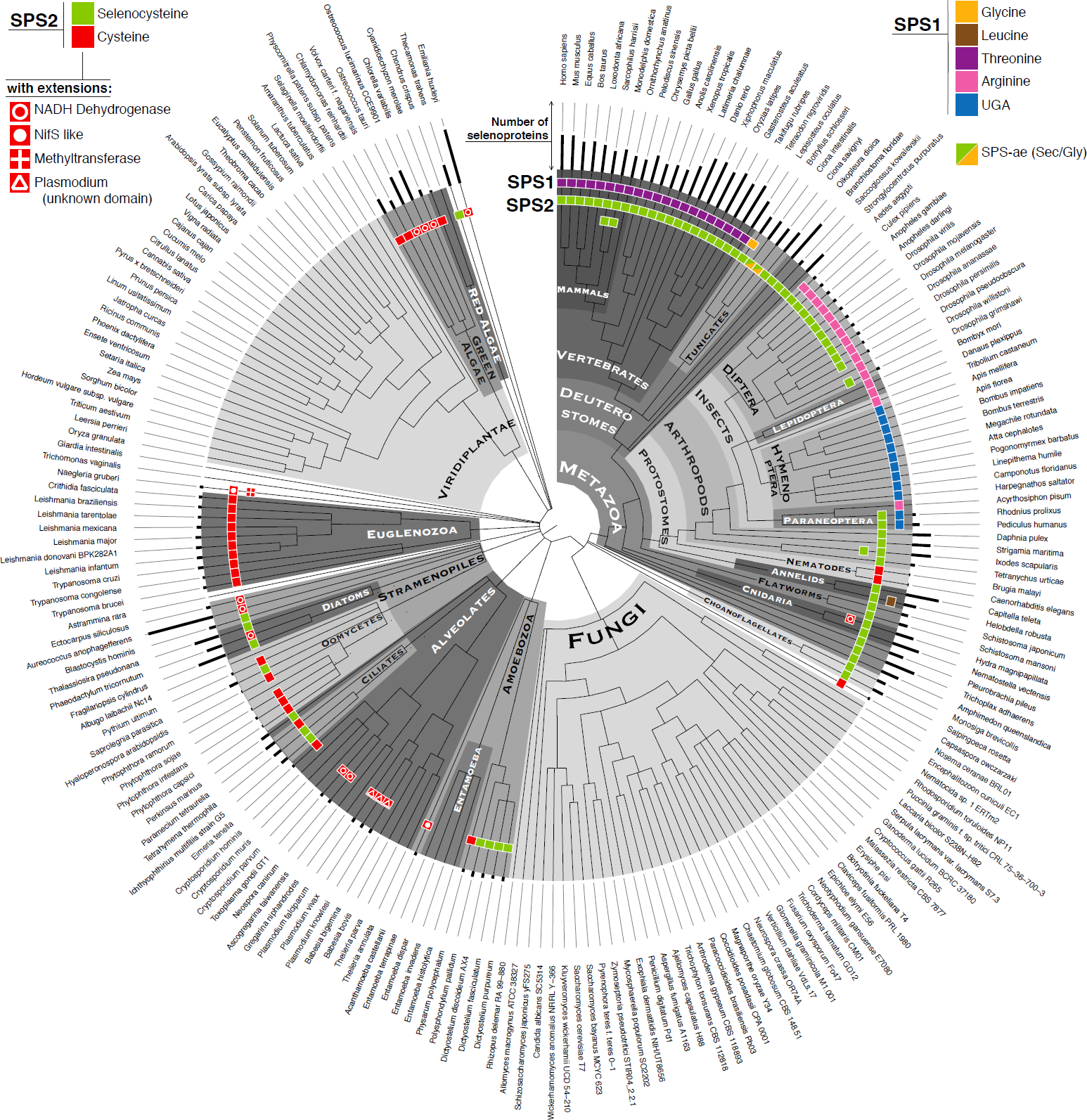
Phylogenetic profile of SPS genes and approximate selenoproteome size of eukaryotes. The plot recapitulates the results on 505 genomes analyzed, summarized to 213 displayed here. The tree (from NCBI taxonomy) is partitioned in lineages, highlighted in grey tones to help visualization. Near the tips of leaves, the presence of SPS proteins is displayed using colored rectangles. Sec (green) and Cys (red) forms correspond to SPS2 (top left legend), while the other homologues to SPS1 (top right legend). The SPS gene extensions found for some Cys-SPS2 are indicated with symbols inside the rectangle (top left legend). The number of selenoproteins predicted in each genome is indicated with a black vertical bar extending from the tips.

Overall, the Sec trait exhibits a rather scattered distribution in protists reflecting a dynamic evolution similar to bacteria. We found SPS2 (and therefore selenoproteins) scattered across *Stramenopiles*, *Alveolata*, *Amoebozoa* and other protist lineages, presumably due to multiple independent events of selenoprotein extinction. In contrast, we found selenoproteins in all investigated *Kinetoplastida* (*Euglenozoa*), including the parasites *Trypanosoma* and *Leishmania* (Lobanov et al. 2006)(Cassago et al. 2006). We did not find any *bona fide* SPS2 nor selenoproteins in Fungi or land plants (*Embryophyta*), despite many genomes searched (284 and 41, respectively, a subset of which are shown in Figure 2). In contrast, green algae genomes contain large numbers of selenoproteins as previously reported (Novoselov et al. 2002)(Palenik et al. 2007). The largest number was in the pelagophyte algae *Aureococcus anophagefferens* (heterokont, Stramenopiles), known for its rich selenoproteome (Gobler et al. 2013). All metazoans encode SPS2 and selenoproteins, with exceptions detected so far only in some insects (Chapple and Guigó 2008)(Lobanov et al. 2008) and some nematodes (Otero et al. 2014).

As in prokaryotes, we found a few protist genomes in which SPS2 is fused to other genes. Fusions with a NADH dehydrogenase–like domain are also the most common, and scattered among many taxa (Figure 2). We detected SPS2-NifS fusions in the amoeba *Acanthamoeba castellani* and the heterolobosean *Naegleria gruberi*. In these two genomes, we found additional SPS2 candidates (Supplementary Material S2). In *N. gruberi*, it is fused to a polypeptide containing a methyltransferase domain (da Silva et al. 2013). Finally, all SPS2 proteins in Plasmodium species were found to possess a large polypeptide extension (>500 amino acids). This domain shows no homology with any known protein, and its function remains unknown. We did not find convincing SPS fusions in non-protist eukaryotes (see Supplementary Material S2).

### Independent duplications of SPS2 generates SPS1 proteins in metazoans

In many metazoan lineages we detected additional SPS genes, which are neither selenoproteins nor Cys-homologues. Since, within metazoans, SPS-Cys are found only in nematodes, and, outside metazoans, additional SPS genes are absent (Figure 2), we argue that the last common metazoan ancestor possessed a single SPS gene with Sec (i.e., it was SPS2). Our results suggest that the additional SPS genes have originated by multiple independent duplications of SPS2. We have specifically identified four independent duplications (Figures 3 and 4). One SPS duplication occurred at the root of the vertebrates, probably as part of one of the reported rounds of whole genome duplications (Dehal and Boore, 2005). Another duplication occurred within tunicates, likely originated by retro-transposition of an alternative isoform of SPS2. Another duplication occurred within annelids, at the root of the *Clitellata* lineage. Finally, a duplication occurred at the root of insects. In each of these duplications, a specific substitution of the Sec residue was fixed: threonine in vertebrates, glycine in tunicates, and leucine in annelids. In insects, however, the UGA codon was maintained after duplication, and substituted in some lineages by arginine. There were at least two independent UGA to arginine substitutions in *Paraneoptera* and *Endopterygota*. Insects without selenoproteins lost the original SPS2 gene, but maintained the duplicated copy.

**Figure 3:**
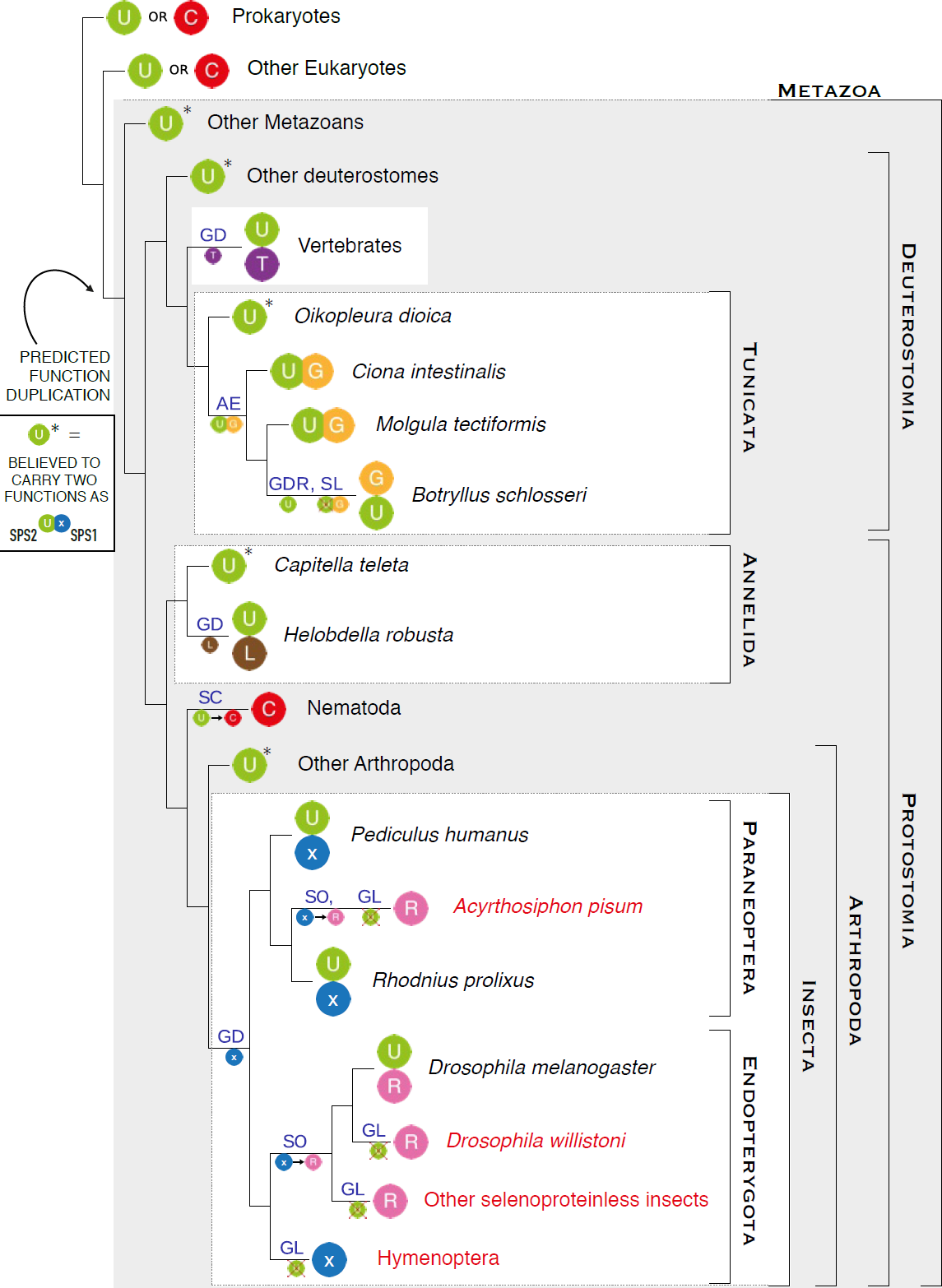
Parallel gene duplications of SPS proteins in metazoan. The plot summarizes the phylogenetic history of metazoan SPS genes, consisting of parallel and convergent events of gene duplication followed by subfunctionalization. Each colored ball represents a SPS gene, indicating the residue found at the UGA or homologous codon (U for selenocysteine, C for cysteine, T for threonine, G for glycine, L for leucine, R for arginine, x for unknown residue). The gene structures are schematically displayed in Figure 4. The names of the insect species lacking selenoproteins are in red. The main genomic events shaping SPS genes are indicated on the branches: GD whole gene duplication, GDR gene duplication by retro-transposition, AE origin of an alternative exon, SL Sec loss, SC conversion of Sec to Cys, SO conversion of Sec to something other than Cys, GL gene loss. In our subfunctionalization hypothesis (see text), we map the origin of a dual function at the root of metazoa.

**Figure 4:**
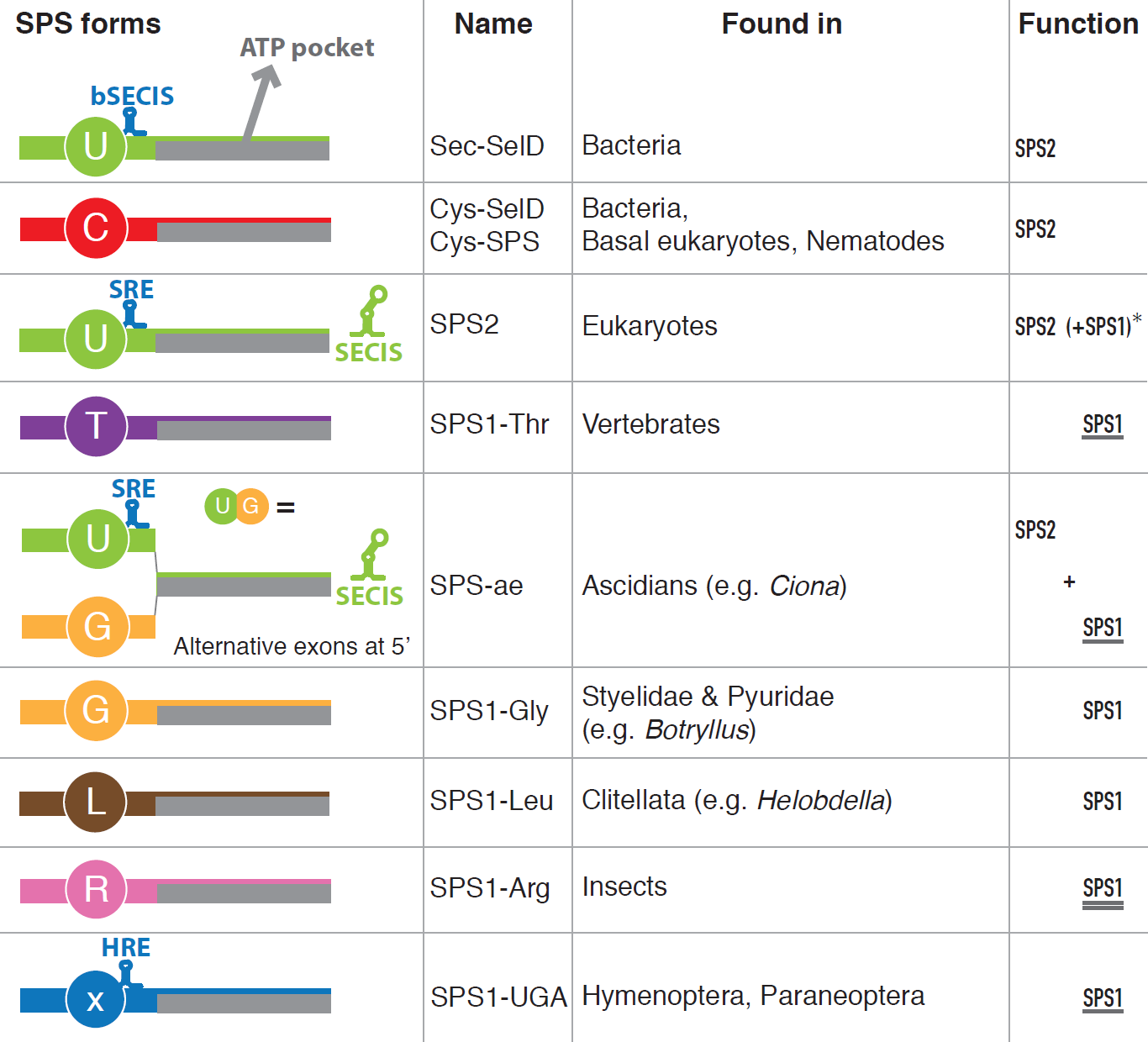
Structure and function of the identified SPS genes. SPS proteins are classified according to the residue found at the UGA or homologous position (see also Figure 3). The presence of specific secondary structures is also indicated: bSECIS for bacterial SECIS element, SRE for Sec recoding element (Howard et al. 2005), SECIS for eukaryotic SECIS element, HRE for hymenopteran readthrough element. The rightmost column indicates the functions predicted for the SPS proteins. SPS2 function is the synthesis of SeP. SPS1 function is defined as the uncharacterized molecular function of *Drosophila* SPS1-Arg (double underlined), which is likely to be similar to that of other SPS1 genes, as suggested by KO-rescue experiments in *Drosophila* (underlined). *: for eukaryotic SPS2, the parentheses indicate that some such genes are predicted to possess both SPS1 and SPS2 functions, those marked also with a star (*) in Figure 3 (basically all metazoans with no SPS1 protein in the same genome).

Therefore, as a universal trend, Cys- or Sec- containing SPS genes (to which we refer as SPS2) are found in all metazoan (eukaryotic) genomes encoding selenoproteins, while eukaryotic genomes containing only SPS genes carrying amino acids other than Sec or Cys at the homologous UGA position do not encode selenoproteins. Thus, the duplicated non-Cys, non-Sec copies of SPS2 in human and fly are unlikely to carry a function related to selenoprotein synthesis. Since in human and fly, they are commonly referred to as SPS1 (Xu et al. 2007b), we will collectively refer to all non-Cys, non-Sec SPS2 duplications in metazoans as SPS1 (e.g. SPS1-Thr for human SPS1).

The SPS duplications had a detectable effect on the selective pressure acting on SPS2. In fact, we consistently detected accelerated rates of non-synonymous vs synonymous substitution in the parental SPS2 gene after each duplication (see Methods and Supplementary Material S3), consistent with a relaxation of the selective constraints acting upon the gene. As a result, SPS1 genes show overall higher sequence similarity across metazoans than SPS2 genes after duplication (80% vs 70%). Also, the predicted ancestral SPS2 genes are more similar to SPS1 genes, than to SPS2 genes after duplication. The higher divergence of SPS2 sequences after duplication appears to be mostly localized in the C-terminal domain (Supplementary Figure SM3.6).

In the next section, we briefly describe each SPS duplication separately.

### SPS phylogeny in vertebrates

All non-vertebrate deuterostomes (except tunicates) and Cyclostomata (jawless vertebrates, such as lampreys) encode only one SPS2 gene, while all Gnasthostomata possess SPS1 in addition. Thus, we conclude that vertebrate SPS1 (SPS1-Thr) originated from a duplication of SPS2 concomitant with conversion of Sec to threonine (Supplementary Material S3) at the root of Gnasthostomata. The conservation of intron positions within the protein sequence is consistent with duplication involving the whole gene structure, and given its phylogenetic position, this is likely to be part of one of the reported rounds of whole genome duplications at the base of vertebrates (Dehal and Boore, 2005). As recently reported (Mariotti et al. 2012), in mammals the SPS2 gene duplicated again, this time by retro-transposition, generating a second SPS2-Sec gene almost identical to the parental, except for the lack of introns. In placental mammals, the intronless SPS2 replaced functionally the parental gene (which was lost), while non-placental mammals still retain the two copies (see, for example, *Monodelphis domestica* in Figure 2).

### SPS phylogeny in tunicates

Tunicates are the closest outgroup to vertebrates (Delsuc et al. 2006), with ascidians (sea squirts) constituting the best-studied and most sequenced lineage. In the ascidian *Ciona*, we identified a single SPS gene with SECIS — the direct descendant of the ancestral metazoan SPS2. Nonetheless, this gene produces two different protein isoforms, deriving from alternative exon structures at the 5’ end (SPS-ae; Supplementary Material S4). One isoform carries Sec (SPS-Sec, corresponding to the SPS2), while the other one, previously unreported, has a glycine instead (SPS-Gly, corresponding to SPS1). We mapped the origin of the SPS-Gly isoform to the root of ascidians, since we did not find it in the non-ascidian tunicate *Oikopleura dioica* (see Figure 5), but both isoforms (SPS-Sec and SPS-Gly) are encoded by the same gene in the ascidian *Molgula tectiformis*. We also found both forms in the recently sequenced ascidian species *Botryllus schlosseri* (Voskoboynik et al. 2013) and *Halocynthia roretzi*, belonging to the sister lineages of *Styelidae* and *Pyuridae*, respectively. However, in these species the two forms mapped to distinct genomic loci, and they correspond therefore to two different genes (see Supplementary Material S4). SPS-Sec is intronless, and contains a SECIS within the 3’UTR. It corresponds, thus, to SPS2. SPS-Gly possesses instead the ancestral intron structure (very similar to *O.dioica* SPS2), and has no SECIS. It corresponds, therefore, to SPS1. Most likely, the ancestral SPS-sec alternative transcript isoform retro-transposed to the genome at the root of *Styelidae* and *Pyuridae*. This generated a copy that soon replaced functionally the SPS-Sec isoform of the parental gene, which as a result specialized in the production only of the SPS-Gly isoform, as both the Sec coding exon and the SECIS element degenerated. This exemplifies an evolutionary scenario, not frequently reported in the literature, in which alternative transcripts precedes gene duplication, providing a possible intermediary step of how a dual-function protein can escape from adaptive conflict (Hittinger and Carroll 2007).

**Figure 5:**
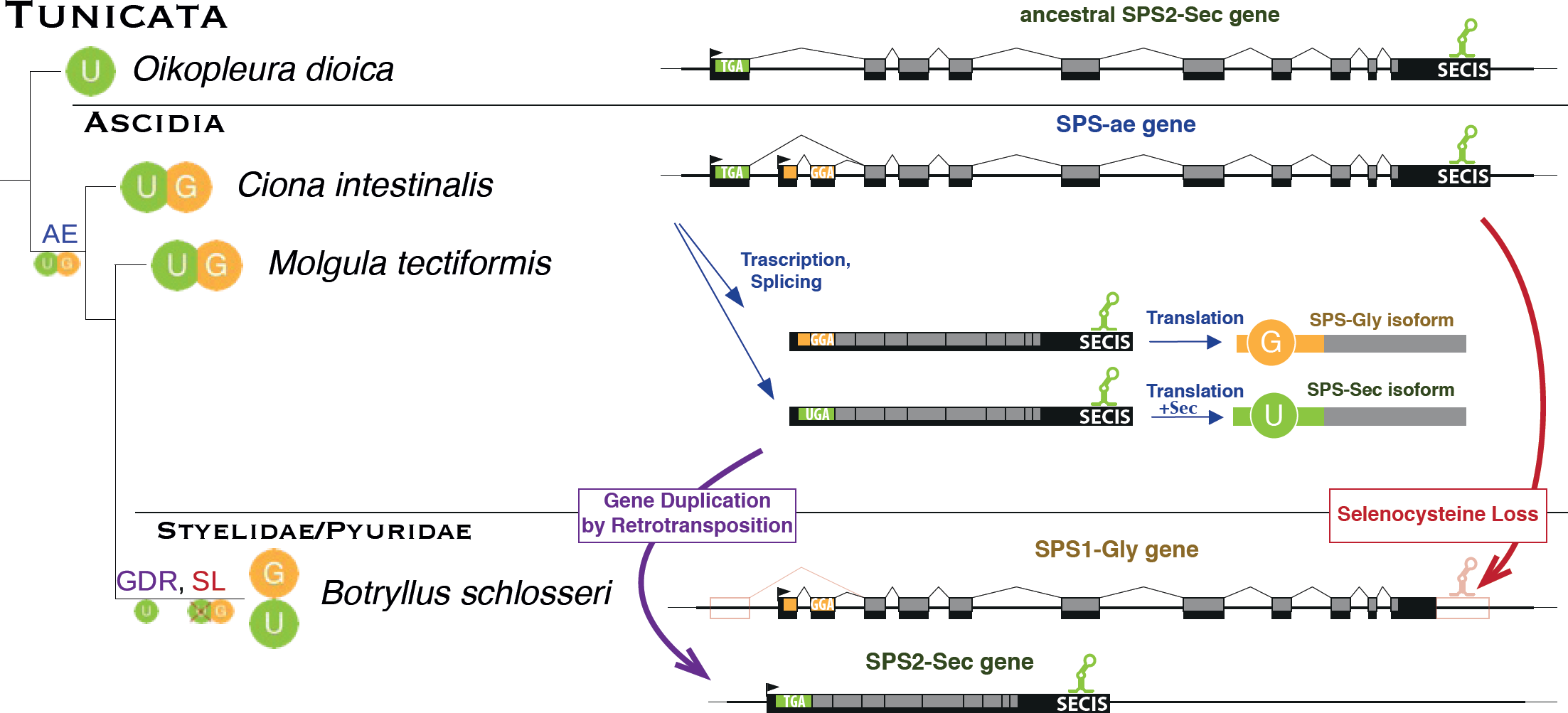
Alternative SPS1/2 transcript isoforms sorted by retrotransposition within ascidians. This figure expands the section for tunicates in Figure 3. At the root of ascidians, the ancestral SPS2-Sec gene acquired a novel SPS1-Gly transcript isoform, through alternative exon usage at the 5’ end (AE). Then, at the root of the ascidian lineage Styelidae and Pyuridae, the SPS2-Sec transcript of this dual SPS1/SPS2 gene (SPS-ae) retrotransposed to the genome creating a novel SPS2-Sec gene (GDR). This presumably triggered the loss of Sec from the parental gene which, as both the SECIS and the UGA containing exon degenerated (SL), specialized only in the production of SPS1-Gly.

### SPS phylogeny in insects

Insects provide a unique framework to study selenoprotein evolution. They have undergone several waves of complete selenoprotein extinction, in which selenoprotein genes were converted to Cys homologues or lost, and the Sec machinery degenerated and/or disappeared. This process occurred in several lineages independently: *Hymenoptera, Lepidoptera, Coleoptera* (or at least *Tribolium castaneum*), and in the *Drosophila willistoni* lineage (Chapple and Guigó 2008)(Lobanov et al. 2008) within Endopterygota, and also in the paraneopteran pea aphid (*Acyrthosiphon pisum* (Aphid-Consortium 2010)). Consistent with its function, the SPS-Sec (SPS2) gene is present in every insect genome coding for selenoproteins, and it is absent in every insect genome not coding for selenoproteins (Figure 2). SPS1 genes, in contrast, are present in all insect genomes. Most insect SPS1 genes (in Lepidoptera, Coleoptera, Diptera, etc) use arginine at the Sec/Cys site (SPS1-Arg, Figures 3 and 4). Instead, SPS1 has a UGA codon at this position in Hymenoptera, and in two non-monophyletic species within paraneopterans: *Rhodnius prolixus* and *Pediculus humanus*. In these, however, and in contrast to hymenopterans, we found an additional SPS-Sec gene with SECIS element (SPS2), as well as, consistently, a number of other selenoproteins and the complete Sec machinery. From all these data (see also Supplementary Material S3), we hypothesize (Figure 3) that all insect SPS1 genes derive from the same SPS2 duplication event occurred approximately at the root of insects, initially generating a UGA-containing, SECIS lacking gene (SPS1-UGA). In most lineages the new gene switched the UGA codon to arginine generating SPS1-Arg proteins. This occurred at least twice independently, in the pea aphid and in the last common ancestor of Coleoptera, Lepidoptera, and Diptera. In Hymenoptera and most Paraneoptera, the gene is still conserved with UGA and no SECIS. The original SPS2 was lost in all lineages where selenoproteins disappeared.

### SPS1-UGA: non-Sec readthrough

The strong conservation of the UGA codon in hymenopteran/paraneopteran SPS1-UGA aligned exactly at the position of the SPS2 Sec-UGA codon is extremely puzzling. SPS1-UGA does not contain a SECIS element. Furthermore, Hymenoptera lack most constituents of the Sec machinery, and these organisms cannot synthesize selenoproteins. However, the striking conservation of the insect SPS1-UGA sequence strongly indicates that is translated and functional. Previously, we had hypothesized that SPS1-UGA could perhaps be translated by a readthrough mechanism not involving Sec insertion (Chapple and Guigó 2008). There is actually a growing evidence for abundant stop codon readthrough in insects, with UGA being the most frequently observed readthrough codon in *Drosophila* (Jungreis et al. 2011).

Here, we have found additional strong evidence in support of the translational recoding of SPS1-UGA. First, recently sequenced hymenopteran genomes (more than ten) all show a clear pattern of protein coding conservation across the UGA, resulting in a readthrough protein of an approximate size of SPS.

Second, we found the hexanucleotide GGG-UG[C/U], which is highly overrepresented next to known viral “leaky” UAG stop codons (Harrell et al. 2002)), to be ultraconserved subsequent to the UGA in SPS1-UGA genes. While the hexanucleotide is found scattered in some other metazoan SPS2 sequences—where it could actually contribute to UGA translation—it is absent from all insect SPS2 genes (Figure 6). Moreover, the hexanucleotide is also absent from insect SPS1 genes having Arg instead of UGA.

**Figure 6:**
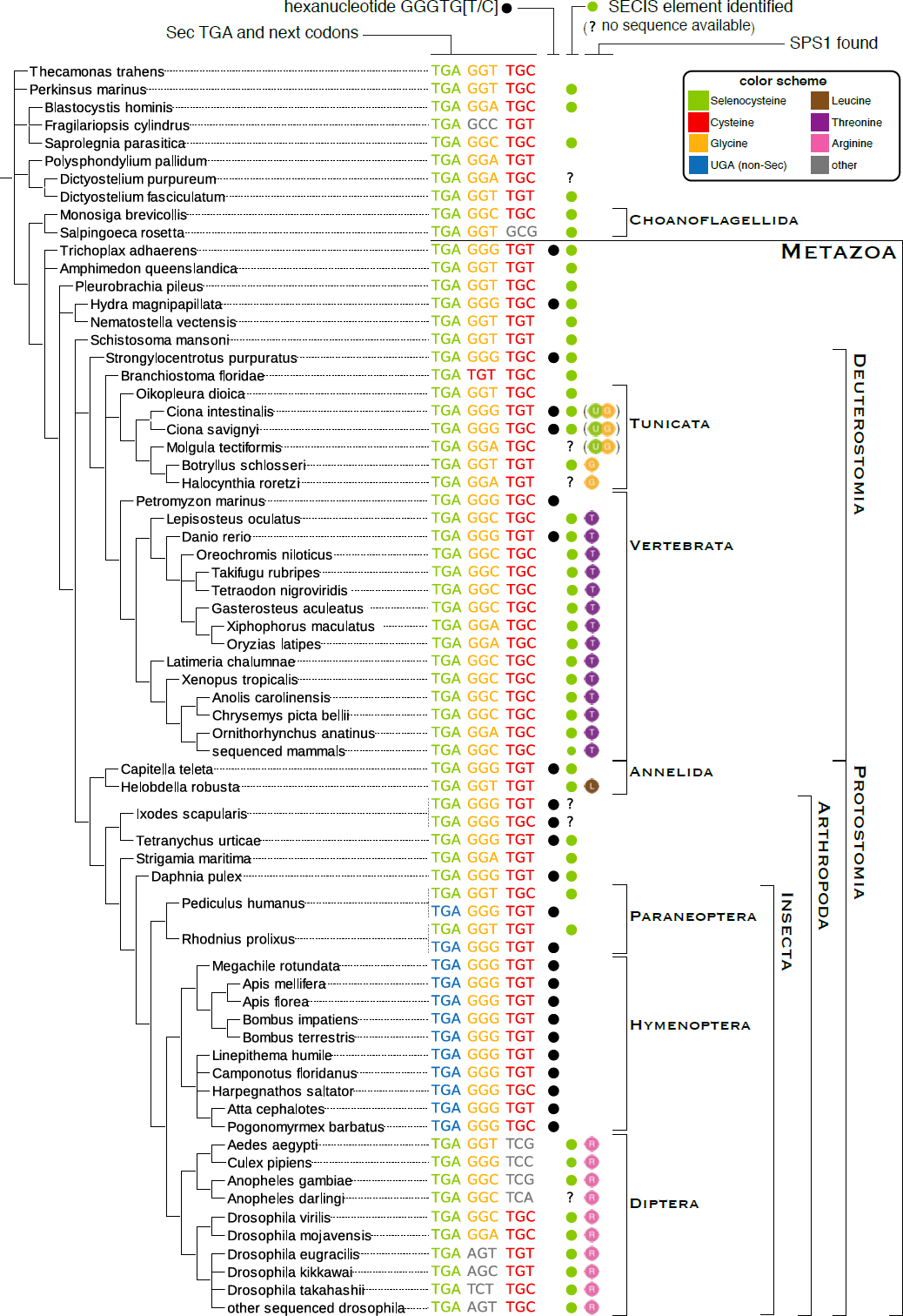
Readthrough-enhancing hexanucleotide in SPS genes. The phylogenetic tree on the left shows the nucleotide sequence alignment at the UGA (or homologous) site in SPS sequences. Only SPS2 and SPS1-UGA genes are shown here. Codons are colored according to their translation, following the same color schema used in Figures 2 and 4 (grey for other amino acids). The presence of the hexanucleotide described in (Harrell et al. 2002) is marked with a black dot. Green dots mark the genes for which a bona-fide SECIS element was identified. The last column indicates the presence of SPS1 genes.

Third, SPS1-UGA contains a conserved secondary structure overlapping the UGA (and therefore the hexanucleotide). We will name this structure HRE, for hymenopteran readthrough element. For its location and structure (Supplementary Material S5, Figure SM5.2), HRE appears to derive from stem-loop structures called SRE (from Sec redefinition elements), previously identified in many selenoprotein genes including SPS2 (Howard et al. 2005), and that promote readthrough activity. Indeed, the SRE element of human SelN gene was functionally characterized (Howard et al. 2005)(Howard et al. 2007), showing that it promotes recoding of UGA codons. In the presence of a downstream SECIS element, Sec insertion is enhanced. In its absence, a Sec-independent readthrough activity is still observed. We hypothesize that HRE of SPS1-UGA has a similar activity. In further support of this, we used RNAz (Gruber et al. 2010) to characterize the secondary structures embedded in the coding sequence of all SPS genes (see Methods and Supplementary Material S5). In prokaryotes, this yielded the bacterial SECIS of the Sec containing SelD genes (Figure SM5.1). In eukaryotes, we obtained stable stem loops (SRE) in the same region of all UGA-containing SPS genes (Figure SM5.2). Strikingly, the largest and most stable structures were in SPS1-UGA, where we predicted HRE as a 3 stem clover-like structure with the UGA in the apex of the middle stem.

Overall, these results strongly suggest that the insect SPS1-UGA gene is translated. Furthermore, a recent proteomics study in the hymenopteran *Cardiocondyla obscurior* (Fuessl et al. 2014) yielded a few peptides mapping to the SPS1-UGA gene—albeit not to the region including the UGA codon. Thus, we cannot unequivocally identify the amino acid that is inserted in response to the UGA codon. Given that we observed two independent UGA to Arg substitutions within insects, Arg could be a potential candidate. Nonetheless, the recognition of a UGA codon by a standard tRNA for Arg would require at best one mismatch in the first position of the codon (third position of the anticodon), which is expected to compromise translation.

### Functional hypothesis: parallel subfunctionalization generates SPS1 proteins

Even though originating from independent gene duplications from the same orthologous gene (Figures 3 and 4), the pattern of strong sequence conservation suggests that SPS1 genes share a common function. That this function is different from SPS2 has been demonstrated for both insect and vertebrates (Persson et al. 1997)(Xu et al. 2007a). We therefore suggest that the ancestral SPS2 protein at the root of metazoans had not only its known catalytic activity (i.e., synthesis of SeP from selenide), but also an additional, secondary function. Eventually, several metazoan lineages split these two, with a new, duplicated protein, SPS1, assuming the secondary function. If this hypothesis is true, then all SPS1 proteins (although paraphyletic) should have similar functions. To test this hypothesis we designed a knockout (KO)-rescue experiment in *Drosophila*. The SPS1 KO mutation (SelD^ptuf^) is lethal in homozygous *Drosophila melanogaster* larvae and results in very reduced and aberrant imaginal disc morphology (Fig. 7A; Alsina et al. 1998). Thus, we investigated if different metazoan SPS1 genes can rescue the phenotype in the *SelD^ptuf^* mutant background. We produced various transgenic flies expressing heterologous SPS1 proteins: human SPS1-Thr, the SPS-Gly isoform from *Ciona intestinalis* SPS-ae, and SPS1-UGA from *Atta cephalotes* (ant, hymenopteran). We then performed crosses to obtain these insertions in flies homozygous for *SelD^ptuf^* (Supplementary Material S6, Figure SM6.1), and examined the morphological phenotypes of third instar larvae wing imaginal discs (Figure 7). We observed significant rescue in the case of *Ciona*, both in size and morphology (Fig. 7C) and partial rescue in size when using the ant SPS1-UGA or the human SPS1-Thr (Fig. 7D, E). These experiments suggest that the tested SPS1 proteins have a similar molecular function, which is, as previously noted, distinct from SeP synthesis.

**Figure 7:**
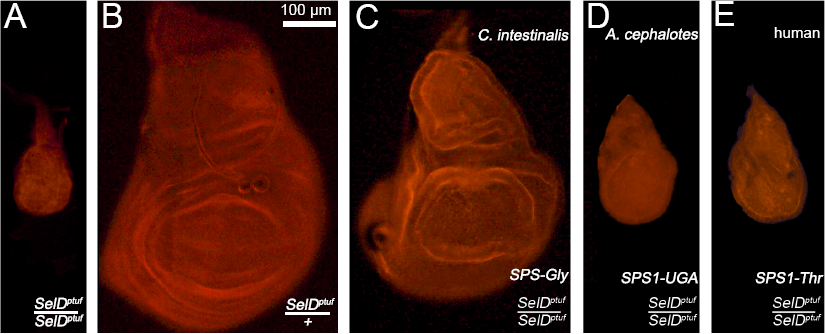
Rescue of the *Drosophila melanogaster* SPS1 null mutant by heterologous SPS1 proteins. All images show wing imaginal discs dissected from larvae with the indicated constructs and genotypes. (A,B) The SPS1 null mutant flies (*SelD^ptuf^* / *SelD^ptuf^*) result in defects in the whole organism, but can be easily monitored in the wing imaginal disc epithelia; the homozygous condition (A) strongly impairs size and morphology, whereas the heterozygous (B) is very similar to the wild type condition. (C-E) The severe homozygous *SelD^ptuf^* / *SelD^ptuf^* phenotype is partially rescued by ubiquitous expression of heterologous SPS1 proteins. (C) *SelD^ptuf^* / *SelD^ptuf^* mutants with *C. intestinalis* SPS1-Gly. (D) *SelD^ptuf^* / *SelD^ptuf^* mutants with *A.cephalotes* SPS1-UGA. (E) *SelD^ptuf^* / *SelD^ptuf^* mutants with human SPS1-Thr.

## Discussion

Gene duplications are generally assumed to be the major evolutionary forces generating new biological functions (Lynch and Conery 2000). However, the mechanisms by means of which duplicated gene copies are maintained and evolve new functions are still poorly understood (Innan and Kondrashov 2010). First, the evolutionary history of genes is inherently difficult to reconstruct, since numerous factors deteriorate and confound phylogenetic signals (Philippe et al. 2011). Second, assigning functions to genes is difficult. Even the very same concept of function is controversial, and there is no universal definition of what constitutes function (Kellis et al. 2014). From a pragmatic standpoint, function is often equated to some sort of biochemical activity, which, in turn, is intimately connected to specific experiments to measure it. But this fails to grasp that genes work in a concerted way within a given biological context, and thus, even similar biochemical activities may result in very different biological functions (phenotypes). Also, experimental validation of function is possible only for a few genes, while for the great majority, function is solely assigned by inference based on sequence similarity. However, there are no universal thresholds to discriminate different functions from levels of sequence divergence, since cases of exist of conserved function with low sequence similarity and conversely, different functions can be carried out by highly similar sequences. For all these reasons, functional evolution of genes is particularly difficult to trace.

Here, we have investigated the evolutionary history of SelD/SPS along the entire tree of life. SPS constitutes a unique gene family to investigate evolution of function. SPS is unique among all genes involved in selenoprotein synthesis in that it is often a selenoprotein itself. Moreover, the UGA-encoded Sec residue is functionally replaceable only with Cys. Therefore, mutations in the UGA to codons other than Cys impair its catalytic function. Consistent with this, we have found that Cys- or Sec- containing SPS genes (which we refer as SPS2) are found in all genomes encoding selenoproteins, while those metazoan genomes containing only SPS genes carrying amino acids other than Sec or Cys at the homologous UGA position do not encode selenoproteins. Our results suggest that the non-Cys, non-Sec SPS genes share a common function, likely unrelated to SeP synthesis, and we refer collectively to them as SPS1. In SPS genes, thus, the amino acid inserted at this single codon position seems to determine unequivocally protein function, and the evolutionary fate of an entire family of proteins. Thanks to this feature, maybe unique among all protein families, we have been able to untangle the complex functional evolution of SPS genes, in particular within metazoans, where it involves a series of independent gene duplications and the associated sub-functionalization events, from an ancestral Sec carrying SPS gene that had both the SPS1 and SPS2 functions. In some ascidians, the duplication originated by selective retro transposition of the SPS2 isoform of a single gene encoding both SPS1 and SPS2 isoforms. Alternative transcript isoforms and gene duplication are considered the main mechanisms contributing to increase protein diversity. They anticorrelate at the genomic scale, so that protein families with many paralogues tend to have fewer transcript variants, and *vice versa* (Talavera et al. 2007). It is therefore expected that some genes shifted from one strategy to the other during evolution: the function of an alternative isoform in one species would be carried by a duplicated copy of the gene in another species. However, other than the ascidian SPS genes reported here, only a handful of cases have been so far reported: the eukaryotic splicing factor *U2AF35* in vertebrates (Pacheco et al. 2004), the *Pax6* gene in *Drosophila* (Dominguez et al. 2004) and the *mitf* gene in fishes (Altschmied et al. 2002).

Gene duplication and alternative transcription differ in the efficiency with which they separate functions. Duplications fully segregate the functions to different loci—allowing both independent sequence evolution and independent regulation. In contrast, alternative transcription/splicing isoforms typically share regions of identical sequence, as well as, generally, proximal and distal regulatory regions. Thus, when a single gene carries a dual function (even if exerted through alternative isoforms), certain sequence segments are under two simultaneous sources of selection, which may be in competition. Gene duplication and subfunctionalization provides a way to entirely escape from the adaptive conflict (Hittinger and Carroll 2007). The fact that convergent subfunctionalization duplications occurred several times independently within metazoans is suggestive of selective advantage in having the two SPS functions carried out by different genes. Indeed, SPS duplications in metazoans partially released the selective pressure acting on SPS2, as we detected accelerated evolution in the parental SPS2 gene after each duplication.

Overall, the history of metazoan SPS genes contributes a striking example how function evolves across orthologs and paralogs through a complex pattern of duplication and loss events (Gabaldón and Koonin 2013). Specifically, it constitutes a prototypical case of a particular “functional evolution path”: an ancestral state of dual/redundant function, followed by subfunctionalization through gene duplication which occurs independently in parallel lineages of descent. Analogous cases have been previously reported (e.g. the β-*catenin/armadillo* gene during insect evolution (Bao et al. 2012)). Parallel gene duplications result in a complex pattern of orthology/paralogy relationship (Gabaldón 2008). Within monophyletic lineages with both SPS1 and SPS2 originated by a duplication event (e.g. vertebrates), these genes are always paralogous, but both genes are phylogenetically co-orthologous to SPS2 in lineages with a single SPS (e.g. flatworms). Also, these genes are phylogenetically co-orthologous to both SPS1 and SPS2 in other lineages with a distinct, parallel duplication event (e.g. insects). Nonetheless, if we assume that sub-functionalization occurred in the same way in the different lineages, then the SPS1 genes generated independently can be considered functional orthologous, despite the lack of direct phylogenetic descent. In this view, it may appear counter-intuitive that SPS1 genes have replaced the Sec/Cys site with lineage-specific amino acids which are so dissimilar. This suggests that the residue occurring at this particular position is irrelevant to SPS1 function.

Our results suggest that readthrough of the SPS UGA codon to incorporate amino acids other than Sec played an important role in metazoan SPS duplication and subfunctionalization. Indeed, we hypothesize that simultaneous to the production of UGA containing transcripts, selenoprotein genes (including SPS2) could produce truncated SECIS-lacking mRNAs, due, for instance to inefficient transcription, or through a regulated process, such as alternative usage of 3’ exons or poly-adenylation sites. In fact, translational regulation is known to play an important role for many selenoproteins (Howard et al. 2013). In at least one case (Selenoprotein S), this is achieved by the regulated inclusion/exclusion of the SECIS element in the mature transcripts through alternative splicing (Bubenik et al. 2013). In such SECIS-lacking truncated transcripts, no Sec insertion will take place, and termination of translation at the UGA codon is the most likely outcome. However, if alternative non-SECIS mediated readthrough mechanisms are present, other amino acids could still be incorporated in response to the UGA codon. If this amino acid is other than Sec or Cys, the resulting protein may not be able to perform its original function (as in SPS2/SPS1), but it may develop a novel one. We speculate that this was the case for the ancestral metazoan SPS gene, in which, possibly, the novel SPS1 function emerged in a secondary, non-Sec isoform maybe produced through regulated SECIS inclusion. The readthrough enhancing stem loop structures around the UGA codon (SRE and HRE) found in SPS genes support this hypothesis. These structures are particularly strong in the SECIS-lacking hymenopteran SPS1-UGA genes, which, in addition, contain the readthrough enhancing hexanucleotide. Then, when the SPS gene duplicates, the SECIS is lost in the SPS1 copy, leading to complete subfunctionalization. The whole process is best seen within insects. Hymenoptera, after the SPS duplication, lost SPS2 as selenoproteins disappeared from the genome, but still conserved SPS1-UGA. Paraneopterans with selenoproteins (*R.prolixus* and *P.humanus*) still maintain the two genes (SPS1 and SPS2) with UGA. In pea aphid and in non-hymenopteran Endopterygota, the in-frame UGA in SPS1-UGA mutated to an Arg codon, becoming the “standard” gene known as *Drosophila* SPS1.

It is tempting to speculate that the SRE/HRE stem loop structures, which are found not only in SPS, but also in some other selenoprotein genes (Howard et al. 2005), are the eukaryotic homologues of the bacterial SECIS element (bSECIS, see Figure SM5.1 and SM5.2). Indeed, the bacterial Sec insertion system is different from its eukaryotic counterpart, both regarding the structure and the localization of the SECIS element. In eukaryotes, the SECIS is characterized by a kink-turn core, and it is located in the 3’ UTR. In bacteria, the bSECIS it is a simple stem-loop structure (lacking the kink-turn motif), located immediately downstream of the Sec-UGA within the coding sequence. bSECIS elements are read by SelB, a Sec-specific elongation factor with a specialized N-terminal domain. In eukaryotes, SECIS elements are bound by SBP2, which recognizes specific structural features mainly around its kink-turn core (Krol 2002).

The evolutionary history of the SECIS elements across the domains of life remains largely unexplored. The assumption that SECIS and bSECIS are phylogenetically homologous structures requires the re-location of the bSECIS to the 3’UTR, concomitant with the radical alteration of its structure—for both of which it is difficult to postulate plausible evolutionary mechanisms. In contrast, SRE/HRE are stem loop structures that localize next to the UGA codon, and resemble much more the bSECIS structure than the SECIS does. In addition to structural similarity, bSECIS and HRE/SRE also share functional similarity, since both disfavor termination at Sec-UGA sites during translation. Thus, we hypothesize that the SRE/HRE structures (at least those in SPS genes) are derived from bSECIS, and that the eukaryotic SECIS is an evolutionary innovation unconnected to the bSECIS. After the emergence of the eukaryotic SECIS system, the ancestral bSECIS function was “downgraded” to helper for Sec insertion (SRE). In ancestral metazoans it was kept under selection to allow both Sec-insertion and non-Sec readthrough, as the non-Sec isoform acquired the SPS1 function. Within insects, after the SPS duplication, the ancestral bSECIS structure remained in one of the duplicated copies (SPS1-UGA), specializing only in non-Sec readthrough. This structure has been conserved in hymenopterans and some paraneopterans, becoming what we named here HRE.

In summary, thanks to the singular feature that the amino acid occurring at a single position serves as binary indicator of gene function, we traced the evolutionary history of SelD/SPS genes with unprecedented detail, providing simultaneously a survey of Se utilization traits across the entire tree of life. In metazoans, the SPS phylogeny constitutes a prototypical case of functional evolution, in which dual function is segregated to different loci through independent gene duplications and subsequent convergent subfunctionalization.

## Methods

### Gene prediction

We performed gene prediction using Selenoprofiles ver. 3.0 (Mariotti and Guigó 2010) (http://big.crg.cat/services/selenoprofiles). This program scans nucleotide sequences to predict genes belonging to given protein families, provided as amino acid sequence alignments (profiles). To ensure maximum sensitivity, two manually curated profiles were used for SPS, one containing sequences from all lineages and another only from prokaryotes. Specificity was controlled through a set of filters applied to predicted candidate genes, which include checking similarity to the profile sequences, and best matches among all known proteins in the NCBI non-redundant (NR) database (AWSI score and tag score – see Selenoprofiles manual). We built our SPS gene datasets (available in Supplementary Material S7) searching a large collection of eukaryotic (505) and prokaryotic genomes (8.263 with a non redundant reference subset of 223), downloaded mainly from NCBI. For eukaryotes and for prokaryotes in the reference set, results were manually inspected and filtered to exclude duplicates, pseudogenes (abundant in vertebrates) and contaminations of the genome assemblies. Eukaryotic SECIS elements were searched using the program SECISearch3 (Mariotti et al. 2013).

We also used Selenoprofiles with profiles derived from protein families that are markers for other selenium utilization markers, ybbB and SelA. We used the same program with a comprehensive collection of selenoprotein families in a semi-automatic procedure to probe the number of selenoproteins per lineage (as those displayed in Figures 1 and 2). tRNAscan ver. 1.23 (Lowe and Eddy 1997) and Aragorn ver. 1.2.28 (Laslett and Canback 2004) were used to search for tRNAsec. We noticed the presence of abundant false positives in prokaryotes, lacking the long extra-arm characteristic of tRNAsec. Thus, we focused most of our analysis on the reference set, inspecting manually candidates and filtering out all such cases.

For ciliates all predictions were manually adjusted, given their non-standard genetic code. In addition to genomes, the NCBI EST database was also used to investigate certain eukaryotic lineages of interest, such as tunicates and annelids (Supplementary Materials S3 and S4).

### Phylogenetic analysis

Alignments were computed using T-Coffee ver. 8.95 (Notredame et al. 2000) and sometimes complemented by Mafft ver. 7.017b (Katoh et al. 2005). To deduce the phylogenetic history of SPS, we combined three types of information: topology of gene trees reconstructed from protein sequences, phylogenetic tree of investigated species, and positions of introns in respect to protein sequence. Gene trees were computed by maximum likelihood as explained in (Mariotti et al. 2012) after (Huerta-Cepas et al. 2011), and visualized using the program ETE ver. 2 (Huerta-Cepas et al. 2010). The approximate phylogenetic tree of investigated species was derived from the NCBI taxonomy database (Sayers et al. 2009), and was refined for insects with data from Aphid-Consortium (2010). Figures 1, 2 and supplementary figure SM1.1 were generated with the program ggsunburst, available at http://genome.crg.es/~didac/ggsunburst/. Relative positions of introns were visualized using selenoprofiles_tree_drawer (available within Selenoprofiles) and ETE.

The SPS phylogenetic history presented here has been deduced using parsimony as the main driving principle. Supplementary Material S3 contains a detail description of the process to solve the phylogeny of eukaryotic SPS. Supplementary Material S4 is dedicated to SPS genes in tunicates.

### Evolutionary analysis

To investigate the evolutionary trajectory of SPS genes with regard to their duplications, we performed sequence analysis on a “summary set” codon alignment, containing representatives for all 4 gene duplications here described (Supplementary Figure SM3.6). To improve the prediction of ancestral sequences, we included one outgroup (unduplicated SPS2 gene) for each duplication event. The codon alignment was trimmed to leave out the N-terminal and C-terminal tails, and to remove any insertion in single sequences. Evolutionary analysis was then performed using the program Pycodeml (Mariotti, unpublished). This package runs CodeML, a program which is part of the PAML package (Yang 2007) to predict the sequence at ancestral nodes, and then computes various indexes of sequence evolution. We used the index “dKaKs” (explained hereafter) to estimate the rate of non-synonymous vs. synonymous codon substitutions after each duplication (Supplementary Figure SM3.6). We computed this metric separately for each duplicated SPS2 gene and for each SPS1 gene, always in reference to their last common ancestor (the predicted sequence prior to duplication). At first, this ancestral sequence is scanned, computing the theoretical effect of every possible single-nucleotide change (synonymous, or non-synonymous substitution). Then, the sequence of the gene under analysis is compared to the ancestral sequence, counting how many of the possible changes were observed. The dKaKs index is computed as the proportion of observed vs. possible non-synonymous substitutions, divided by the proportion of observed vs. possible synonymous substitutions. A small dKaKs indicates strong purifying selection acting on the protein sequence; a value of 1.0 indicates neutral evolution (no constraints). sec">Supplementary Material S3 contains the results of this analysis.

### Detection of extensions and fusions

We used two different strategies to detect additional domains in SPS genes. First, we searched for annotated SPS fusions. We ran our SPS gene set with BLASTp (Altschul et al. 1997) against the NCBI NR database. Then we parsed the results, searching for large stretches of sequence of a matched NR protein that were not included in the output BLAST alignment. Second, we looked in genomes for any SPS extension. We expanded each predicted SPS gene at both sides until a stop codon was reached, and we ran the extensions with BLASTp against NCBI NR database. All candidates from the two methods were merged, clustered by similarity, and manually inspected. Conservation in multiple species was used as criteria to exclude artifacts possibly caused by our detection method or by imperfect genome assemblies. For the most interesting cases, a new alignment profile was built including the sequence of SPS and of the additional domain, and used to search again the genome sequences. Supplementary Material S2 contains a description of results.

### Prediction of conserved secondary structures

The program RNAz ver. 2.1 (Gruber et al. 2010) was used to predict conserved secondary structures embedded in SPS coding sequences. Initially, we produced a “master alignment” that included the coding sequences of all SPS genes in our dataset. The nucleotide sequence alignments employed were based on the alignment of the corresponding amino acid sequences. Then, the master alignment was used to extract a multitude of “subset alignments”, which included only genes in specific lineages, and/or only specific types of SPS (based on the residue found at the Sec position). RNAz was run either directly on these subset alignments, or instead the program TrimAl ver. 1.4 (Capella-Gutiérrez et al. 2009) was used in advance to reduce the number of sequences. All secondary structures predicted in this way were then manually inspected. For the best candidates, images of consensus structures were generated using the Vienna RNA package (Lorenz et al. 2011)(see Figures SM5.1 and SM5.2). Supplementary Material S5 contains a detailed description of the procedure and of the results, including the list of subset alignments considered.

### Rescue experiments in *Drosophila*

We obtained cDNA for human SPS1 from the Harvard resource core (http://plasmid.med.harvard.edu/PLASMID/). For SPS-ae of *C.intestinalis*, we obtained the cDNA corresponding to the Gly isoform by performing targeted PCR on larvae extracts. We obtained cDNA for SPS1-UGA from *A.cephalotes* by performing targeted PCR on extracts provided by James F.A. Traniello. These cDNAs were cloned through the Gibson method (Gibson et al. 2009) into the *pUAST-attB* vector linearized by double digestion with BglII and XhoI and clones verified by Sanger sequencing. Primers used for cloning are reported in Supplementary Material S6. Transgenic flies were obtained following the method described by (Bischof et al. 2007). Line y*w M{e.vas-int.DM}ZH-2A; 3: M{RFP.attP}ZH-86Fb* was used to direct the insertion into the 3R chromosome (86F). Crosses were designed to obtain expression of each transgenic SPS1 into a SelD^ptuf^ homozygous mutant background (Supplementary Material S6, Figure SM6.1). We used the arm-Gal4 line as a driver to activate the *UAS*-cDNA inserts. The final cross was: *SelD^ptuf^*/CyOdfYFP; *UAS*-cDNA insert/MKRS x *SelD^ptuf^*/CyOdfYFP; *arm-Gal4*/TM6B. Imaginal wing discs from third instar larvae were dissected in PBS and stained with Rhodamine Phalloidin from Molecular Probes® (cat #R415).

## Acknowledgements

We thank James F.A. Traniello and Ysabel Milton Giraldo (Boston University, Boston) for providing extracts of *A.cephalotes*. We thank the Drosophila Injection Platform of the Consolider Project (CBM-SO, Madrid) for production of transgenic flies. RG group research was funded by grants BIO2011-26205 from the Spanish Ministry of Science and grant SGR-1430 from the Catalan Government. MM received a FPU doctoral fellowship AP2008-04334 from the Spanish Ministry of Education. TG group research was funded in part by a grant from the Spanish Ministry of Economy and Competitiveness (BIO2012-37161), a grant from the Qatar National Research Fund grant (NPRP 5-298-3-086), and a grant from the European Research Council under the European Union’s Seventh Framework Programme (FP/2007-2013) / ERC (Grant Agreement n. ERC-2012-StG-310325). VNG group research was supported by NIH GM061603.

## Supplementary Material

Find next the following supplementary sections:

- S1: SelD in prokaryotes
- S2: Gene fusions and extensions
- S3: Phylogeny of eukaryotic SPS proteins
- S4: Alternative isoforms sorted by gene duplication in ascidians
- S5: Secondary structures within coding sequences of SPS genes
- S6: Rescue experiments in *Drosophila*
- S7: Datasets

## Abbreviations used

Sec: selenocysteine
Se: selenium
SeU: selenouridine
SeP: selenophosphate
SPS: selenophosphate synthetase
SelD: selenophosphate synthetase (prokaryotes)
SelA: selenocysteine synthase (prokaryotes)
ybbB: selenouridine synthase (prokaryotes)
Cys: cysteine
Arg: arginine
Thr: threonine
Gly: glycine
Leu: leucine

## References

Allmang C, Wurth L, Krol A. 2009. The selenium to selenoprotein pathway in eukaryotes: more molecular partners than anticipated. Biochim Biophys Acta, 1790(11):1415–23.

Alsina B, Serras F, Baguna J, Corominas M. 1998. patufet, the gene encoding the Drosophila melanogaster homologue of selenophosphate synthetase, is involved in imaginal disc morphogenesis. Mol Gen Genet, 257(2):113–123.

Altschmied J, Delfgaauw J, Wilde B, Duschl J, Bouneau L, Volff JN, Schartl M. 2002. Subfunctionalization of duplicate mitf genes associated with differential degeneration of alternative exons in fish. Genetics, 161(1):259–67.

Altschul S, Madden T, Schaffer A, Zhang J, Zhang Z, Miller W, Lipman D. 1997. Gapped BLAST and PSI-BLAST: a new generation of protein database search programs. Nucleic acids res, 25(17):3389.

Aphid-Consortium (The International Aphid Genomics Consortium). 2010. Genome sequence of the pea aphid Acyrthosiphon pisum. PLoS Biol, 8(2):e1000313.

Bao R, Fischer T, Bolognesi R, Brown SJ, Friedrich M. 2012. Parallel duplication and partial subfunctionalization of β-catenin/armadillo during insect evolution. Mol Biol Evol, 29(2):647–62.

Bischof J, Maeda RK, Hediger M, Karch F, Basler K. 2007. An optimized transgenesis system for Drosophila using germ-line-specific phiC31 integrases. Proc Natl Acad Sci U S A, 104(9):3312–7.

Bubenik JL, Miniard AC, Driscoll DM. 2013. Alternative transcripts and 3’UTR elements govern the incorporation of selenocysteine into selenoprotein S. PLoS One, 8(4):e62102.

Capella-Gutiérrez S, Silla-Martínez JM, Gabaldón T. 2009. trimAl: a tool for automated alignment trimming in large-scale phylogenetic analyses. Bioinformatics, 25(15):1972–3.

Cassago A, Rodrigues EM, Prieto EL, Gaston KW, Alfonzo JD, Iribar MP, Berry MJ, Cruz AK, Thiemann OH. 2006. Identification of Leishmania selenoproteins and SECIS element. Mol Biochem Parasitol, 149(2):128–34.

Castellano S, Andrés AM, Bosch E, Bayes M, Guigó R, Clark AG. 2009. Low exchangeability of selenocysteine, the 21st amino acid, in vertebrate proteins. Mol Biol Evol, 26(9):2031–40.

Chapple CE, Guigó R. 2008. Relaxation of selective constraints causes independent selenoprotein extinction in insect genomes. PLoS One, 13;3(8):e2968.

da Silva MTA, Caldas VEA, Costa FC, Silvestre DAMM, Thiemann OH. 2013. Selenocysteine biosynthesis and insertion machinery in Naegleria gruberi. Mol Biochem Parasitol, 188(2):87–90.

Dehal P, Boore JL. 2005. Two rounds of whole genome duplication in the ancestral vertebrate. PLoS Biol, 3(10):e314.

Delsuc F, Brinkmann H, Chourrout D, Philippe H. 2006. Tunicates and not cephalochordates are the closest living relatives of vertebrates. Nature, 439(7079):965–8.

Dominguez M, Ferres-Marco D, Gutierrez-Aviño FJ, Speicher SA, Beneyto M. 2004. Growth and specification of the eye are controlled independently by Eyegone and Eyeless in Drosophila melanogaster. Nat Genet, 36(1):31–9.

Fuessl M, Reinders J, Oefner PJ, Heinze J, Schrempf A. 2014. Selenophosphate synthetase in the male accessory glands of an insect without selenoproteins. J Insect Physiol, 2014 Dec;7146–51.

Gabaldón T. 2008. Large-scale assignment of orthology: back to phylogenetics?. Genome Biol, 30;9(10):235.

Gabaldón T, Koonin EV. 2013. Functional and evolutionary implications of gene orthology. Nat Rev Genet, 14(5):360–6.

Gibson DG, Young L, Chuang RY, Venter JC, Hutchison CA3rd, Smith HO. 2009. Enzymatic assembly of DNA molecules up to several hundred kilobases. Nat Methods, 6(5):343–5.

Gobler CJ, Lobanov AV, Tang Y-Z, Turanov AA, Zhang Y, Doblin M, Taylor GT, Sañudo Wilhelmy SA, Grigoriev IV, Gladyshev VN. 2013. The central role of selenium in the biochemistry and ecology of the harmful pelagophyte, Aureococcus anophagefferens. ISME J, 7(7):1333–43.

Gruber AR, Findeiß S, Washietl S, Hofacker IL, Stadler PF. 2010. RNAz 2.0: improved noncoding RNA detection. Pac Symp Biocomput, 69–79.

Haft DH, Self WT. 2008. Orphan SelD proteins and selenium-dependent molybdenum hydroxylases. Biol Direct, 34.

Harrell L, Melcher U, Atkins JF. 2002. Predominance of six different hexanucleotide recoding signals 3’ of read-through stop codons. Nucleic Acids Res, 30(9):2011–2017.

Hittinger CT, Carroll SB. 2007. Gene duplication and the adaptive evolution of a classic genetic switch. Nature, 449(7163):677–81.

Howard MT, Aggarwal G, Anderson CB, Khatri S, Flanigan KM, Atkins JF. 2005. Recoding elements located adjacent to a subset of eukaryal selenocysteine-specifying UGA codons. EMBO J, 24(8):1596–607.

Howard MT, Carlson BA, Anderson CB, Hatfield DL. 2013. Translational redefinition of UGA codons is regulated by selenium availability. J Biol Chem, 288(27):19401–13.

Howard MT, Moyle MW, Aggarwal G, Carlson BA, Anderson CB. 2007. A recoding element that stimulates decoding of UGA codons by Sec tRNA[Ser]Sec. RNA, 13(6):912–20.

Huerta-Cepas J, Capella-Gutierrez S, Pryszcz LP, Denisov I, Kormes D, Marcet-Houben M, Gabaldón T. 2011. PhylomeDB v3.0: an expanding repository of genome-wide collections of trees, alignments and phylogeny-based orthology and paralogy predictions. Nucleic Acids Res, 39(Database issue):D556–60.

Huerta-Cepas J, Dopazo J, Gabaldón T. 2010. ETE: a python Environment for Tree Exploration. BMC Bioinformatics, 1124.

Innan H, Kondrashov F. 2010. The evolution of gene duplications: classifying and distinguishing between models. Nat Rev Genet, 11(2):97–108.

Jiang L, Ni J, Liu Q. 2012. Evolution of selenoproteins in the metazoan. BMC Genomics, 13446.

Jungreis I, Lin MF, Spokony R, Chan CS, Negre N, Victorsen A, White KP, Kellis M. 2011. Evidence of abundant stop codon readthrough in Drosophila and other metazoa. Genome Res, 21(12):2096–113.

Katoh K, Kuma K-i, Toh H, Miyata T. 2005. MAFFT version 5: improvement in accuracy of multiple sequence alignment. Nucleic Acids Res, 33(2):511–8.

Kellis M, Wold B, Snyder MP, Bernstein BE, Kundaje A, Marinov GK, Ward LD, Birney E, Crawford GE, Dekker J et al. 2014. Defining functional DNA elements in the human genome. Proc Natl Acad Sci U S A, 111(17):6131–8.

Kim IY, Guimarães MJ, Zlotnik A, Bazan JF, Stadtman TC. 1997. Fetal mouse selenophosphate synthetase 2 (SPS2): characterization of the cysteine mutant form overproduced in a baculovirus-insect cell system. Proc Natl Acad Sci U S A, 94(2):418–21.

Krol A. 2002. Evolutionarily different RNA motifs and RNA-protein complexes to achieve selenoprotein synthesis. Biochimie, 84(8):765–774.

Kryukov G, Gladyshev V. 2004. The prokaryotic selenoproteome. EMBO Rep, 5(5):538.

Laslett D, Canback B. 2004. ARAGORN, a program to detect tRNA genes and tmRNA genes in nucleotide sequences. Nucleic Acids Res, 32(1):11–6.

Lacourciere GM, Mihara H, Kurihara T, Esaki N, Stadtman TC. 2000. Escherichia coli NifS-like proteins provide selenium in the pathway for the biosynthesis of selenophosphate. J Biol Chem, Aug 4;275(31):23769–73.

Lee KH, Shim MS, Kim JY, Jung HK, Lee E, Carlson BA, Xu X-M, Park JM, Hatfield DL, Park T et al. 2011. Drosophila selenophosphate synthetase 1 regulates vitamin B6 metabolism: prediction and confirmation. BMC Genomics, 12426.

Lobanov AV, Gromer S, Salinas G, Gladyshev VN. 2006. Selenium metabolism in Trypanosoma: characterization of selenoproteomes and identification of a Kinetoplastida-specific selenoprotein. Nucleic Acids Res, 34(14):4012.

Lobanov AV, Fomenko DE, Zhang Y, Sengupta A, Hatfield DL, Gladyshev VN. 2007. Evolutionary dynamics of eukaryotic selenoproteomes: large selenoproteomes may associate with aquatic life and small with terrestrial life. Genome Biol, 8(9):R198.

Lobanov AV, Hatfield DL, Gladyshev VN. 2008. Selenoproteinless animals: selenophosphate synthetase SPS1 functions in a pathway unrelated to selenocysteine biosynthesis. Protein Sci, 17(1):176.

Lobanov AV, Hatfield DL, Gladyshev VN. 2009. Eukaryotic selenoproteins and selenoproteomes. Biochim Biophys Acta, 1790(11):1424–8.

Lorenz R, Bernhart SH, Höner zu Siederdissen C, Tafer H, Flamm C, Stadler PF, Hofacker IL. 2011. ViennaRNA Package 2.0. Algorithms Mol Biol, 24; 6:26.

Lowe T, Eddy S. 1997. tRNAscan-SE: A program for improved detection of transfer RNA genes in genomic sequence. Nucleic Acids Res, 25(5):955.

Lynch M, Conery JS. 2000. The evolutionary fate and consequences of duplicate genes. Science, 290(5494):1151–5.

Mariotti M, Guigó R. 2010. Selenoprofiles: profile-based scanning of eukaryotic genome sequences for selenoprotein genes. Bioinformatics, 26(21):2656–63.

Mariotti M, Ridge PG, Zhang Y, Lobanov AV, Pringle TH, Guigó R, Hatfield DL, Gladyshev VN. 2012. Composition and evolution of the vertebrate and mammalian selenoproteomes. PLoS One, 7(3):e33066.

Mariotti M, Lobanov AV, Guigo R, Gladyshev VN. 2013. SECISearch3 and Seblastian: new tools for prediction of SECIS elements and selenoproteins. Nucleic Acids Res, 41(15):e149.

Morey M, Serras F, Corominas M. 2003. Halving the selenophosphate synthetase gene dose confers hypersensitivity to oxidative stress in Drosophila melanogaster. FEBS Lett, 534(1-3):111–4.

Notredame C, Higgins DG, Heringa J. 2000. T-Coffee: A novel method for fast and accurate multiple sequence alignment. J Mol Biol, 302(1):205–17.

Novoselov SV, Rao M, Onoshko NV, Zhi H, Kryukov GV, Xiang Y, Weeks DP, Hatfield DL, Gladyshev VN. 2002. Selenoproteins and selenocysteine insertion system in the model plant cell system, Chlamydomonas reinhardtii. EMBO J, 21(14):3681.

Otero L, Romanelli-Cedrez L, Turanov AA, Gladyshev VN, Miranda-Vizuete A, Salinas G. 2014. Adjustments, extinction, and remains of selenocysteine incorporation machinery in the nematode lineage. RNA, 20(7):1023–34.

Pacheco TR, Gomes AQ, Barbosa-Morais NL, Benes V, Ansorge W, Wollerton M, Smith CW, Valcárcel J, Carmo-Fonseca M. 2004. Diversity of vertebrate splicing factor U2AF35: identification of alternatively spliced U2AF1 mRNAS. J Biol Chem, 279(26):27039–49.

Palenik B, Grimwood J, Aerts A, Rouzé P, Salamov A, Putnam N, Dupont C, Jorgensen R, Derelle E, Rombauts S et al. 2007. The tiny eukaryote Ostreococcus provides genomic insights into the paradox of plankton speciation. Proc Natl Acad Sci U S A, 104(18):7705–10.

Palioura S, Sherrer RL, Steitz TA, Söll D, Simonovic M. 2009. The human SepSecS-tRNASec complex reveals the mechanism of selenocysteine formation. Science, 325(5938):321–325.

Persson BC, Böck A, Jäckle H, Vorbrüggen G. 1997. SelD homolog from Drosophila lacking selenide-dependent monoselenophosphate synthetase activity. J Mol Biol, 274(2):174–80.

Philippe H, Brinkmann H, Lavrov DV, Littlewood DT, Manuel M, Wörheide G, Baurain D. 2011. Resolving difficult phylogenetic questions: why more sequences are not enough. PLoS Biol, 9(3):e1000602.

Pruitt KD, Tatusova T, Brown GR, Maglott DR. 2012. NCBI Reference Sequences (RefSeq): current status, new features and genome annotation policy. Nucleic Acids Res, 40(Database issue):D130–5.

Romero H, Zhang Y, Gladyshev VN, Salinas G. 2005. Evolution of selenium utilization traits. Genome Biol, 6(8):R66.

Rother M, Resch A, Wilting R, Böck A. 2001. Selenoprotein synthesis in archaea. Biofactors, 14(1-4):75–83.

Sayers EW, Barrett T, Benson DA, Bolton E, Bryant SH, Canese K, Chetvernin V, Church DM, Dicuccio M, Federhen S et al. 2009. Database resources of the National Center for Biotechnology Information. Nucleic Acids Res, 38(Database issue)D5–16.

Small-Howard A, Morozova N, Stoytcheva Z, Forry EP, Mansell JB, Harney JW, Carlson BA, Xu X-M, Hatfield DL, Berry MJ. 2006. Supramolecular complexes mediate selenocysteine incorporation in vivo. Mol Cell Biol, 26(6):2337–46.

Squires J, Berry M. 2008. Eukaryotic Selenoprotein Synthesis: Mechanistic Insight Incorporating New Factors and New Functions for Old Factors. IUBMB Life, 60(4):232–235.

Srivastava M, Mallard C, Barke T, Hancock LE, Self WT. 2011. A selenium-dependent xanthine dehydrogenase triggers biofilm proliferation in Enterococcus faecalis through oxidant production. J Bacteriol, 193(7):1643–52.

Stock T, Rother M. 2009. Selenoproteins in Archaea and Gram-positive bacteria. Biochimica Biophysica Acta, 1790(11):1520–1532.

Su D, Ojo TT, Söll D, Hohn MJ. 2012. Selenomodification of tRNA in archaea requires a bipartite rhodanese enzyme. FEBS Lett, 586(6):717–21.

Talavera D, Vogel C, Orozco M, Teichmann SA, de la Cruz X. 2007. The (in)dependence of alternative splicing and gene duplication. PLoS Comput Biol, 3(3):e33.

Tamura T, Yamamoto S, Takahata M, Sakaguchi H, Tanaka H, Stadtman TC, Inagaki K. 2004. Selenophosphate synthetase genes from lung adenocarcinoma cells: Sps1 for recycling L-selenocysteine and Sps2 for selenite assimilation. Proc Natl Acad Sci U S A, 101(46):16162–7.

Taskov K, Chapple C, Kryukov GV, Castellano S, Lobanov AV, Korotkov KV, Guigó R, Gladyshev VN. 2005. Nematode selenoproteome: the use of the selenocysteine insertion system to decode one codon in an animal genome?. Nucleic Acids Res, 33(7):2227–38.

Voskoboynik A, Neff NF, Sahoo D, Newman AM, Pushkarev D, Koh W, Passarelli B, Fan HC, Mantalas GL, Palmeri KJ et al. 2013. The genome sequence of the colonial chordate, Botryllus schlosseri. eLife, 2:e00569.

Xu X, Carlson B, Mix H, Zhang Y, Saira K, Glass R, Berry M, Gladyshev V, Hatfield D. 2007a. Biosynthesis of selenocysteine on its tRNA in eukaryotes. PLoS Biol, 5(1):e4.

Xu X, Carlson B, Irons R, Mix H, Zhong N, Gladyshev V, Hatfield D. 2007b. Selenophosphate synthetase 2 is essential for selenoprotein biosynthesis. Biochem J, 404(Pt 1):115.

Yang Z. 2007. PAML 4: phylogenetic analysis by maximum likelihood. Mol Biol Evol, Aug;24(8):1586–91.

Yoshizawa S, Böck A. 2009. The many levels of control on bacterial selenoprotein synthesis. Biochim Biophys Acta, 1790(11):1404–1414.

Zhang Y, Romero H, Salinas G, Gladyshev V. 2006. Dynamic evolution of selenocysteine utilization in bacteria: a balance between selenoprotein loss and evolution of selenocysteine from redox active cysteine residues. Genome Biol, 7(10):R94.

Zhang Y, Turanov A, Hatfield D, Gladyshev V. 2008. In silico identification of genes involved in selenium metabolism: evidence for a third selenium utilization trait. BMC Genomics, 9(1):251.

Zhang Y, Gladyshev VN. 2008. Trends in selenium utilization in marine microbial world revealed through the analysis of the global ocean sampling (GOS) project. PLoS Genet, 4(6):e1000095.

Zhang Y, Gladyshev VN. 2010. General trends in trace element utilization revealed by comparative genomic analyses of Co, Cu, Mo, Ni, and Se. J Biol Chem, 285(5):3393–405.

